# IL-15 superagonist N-803 enhances IFNγ production and alters the trafficking of MAIT cells in SIV+ macaques

**DOI:** 10.1101/2022.06.28.498052

**Authors:** Amy L. Ellis-Connell, Alexis J. Balgeman, Nadean M. Kannal, Karigynn Hansen Chaimson, Anna Batchenkova, Jeffrey T. Safrit, Shelby L. O’Connor

## Abstract

Mucosal Associated Invariant T cells (MAIT cells) are innate T cells that recognize bacterial metabolites and secrete cytokines and cytolytic enzymes to destroy infected target cells. This makes MAIT cells promising targets for immunotherapy to combat bacterial infections. Here, we analyzed the effects of an immunotherapeutic agent, the IL-15 superagonist N-803, on MAIT cell activation, trafficking, and cytolytic function in macaques. We found that N-803 could activate MAIT cells *in vitro* and increase their ability to produce IFNγ in response to bacterial stimulation. To expand upon this, we examined the phenotypes and function of MAIT cells present in samples collected from PBMC, airways (BAL), and lymph nodes (LN) from rhesus macaques that were treated *in vivo* with N-803. N-803 treatment led to a transient 6-7 fold decrease in the total number of MAIT cells in the peripheral blood relative to pre N-803 timepoints. Concurrent with the decrease in cells in the peripheral blood, we observed a rapid decline in the frequency of CXCR3+CCR6+ MAITs. This corresponded with an increase in the frequency of CCR6+ MAITs in BAL, and higher frequencies of ki-67+ and granzyme B+ MAITs in blood, LN, and BAL. Finally, N-803 improved the ability of MAIT cells collected from PBMC and airways to produce IFNγ in response to bacterial stimulation. Overall, N-803 shows the potential to transiently alter the trafficking and enhance the antibacterial activity of MAIT cells which could be combined with other strategies to combat bacterial infections.

## Introduction

Mucosal Associated Invariant T cells (MAIT cells) are a specialized type of innate T cells whose T cell receptor (TCR) can detect bacterial metabolites presented by the MHC-class I related (MR1) molecules on antigen-presenting cells (1, 2, 3). Unlike conventional T cells, crosslinking of the TCR on MAIT cells immediately activates them to perform effector functions, such as secretion of cytokines and cytolytic enzymes (Reviewed in (4)).

Several studies have shown that MAIT cells can recognize bacterial metabolites that are synthesized as part of the bacterial riboflavin biosynthetic pathway (5, 6). Several bacteria produce these metabolites, including *Mycobacterium tuberculosis* (Mtb) (7). MAIT cells have been championed as potentially attractive targets for immunotherapeutic interventions to combat bacterial infections such as Mtb (8). Data supporting an effective role for MAIT cells to combat infectious pathogens is unfortunately mixed. In humans, activated MAIT cells were detected during Mtb infection, and they may be recruited to the lungs (9–11). However, macaque and mouse models are less supportive. There is limited recruitment of MAIT cells to the lungs and they are poorly activated during acute infection with Mtb (12–14). Prophylactic treatment of mice with the riboflavin derivative 5-OP-RU expanded MAIT cells significantly *in vivo*, but those cells did not prevent acute infection with Mtb (15, 16). However, therapeutic vaccination with 5-OP-RU during chronic Mtb infection did reduce bacterial burden (16). Together, these studies imply that MAITs are an intriguing population of cells, but their specific role in antibacterial immunity needs to be more clearly defined.

MAIT cells can also be activated in a TCR-independent manner (17, 18). Cytokines such as IL-7, IL-12, IL-18, and IL-15 can also activate MAIT cells (19–21), leading to increases in cytokine and cytolytic granule production (19–21).

The IL-15 superagonist N-803 has recently gained enthusiasm as an immunotherapeutic agent for combating several cancers and, possibly, HIV (22–26). N-803 contains a constitutively active (N72D) IL-15 molecule and a human IgG Fc receptor that led to improved function and a longer half-life, *in vivo* (27–30). This agent expands CD8 T cells and NK cells (28, 31), increases their cytolytic functions, and improves their trafficking to sites of inflammation and infection (23, 32). N-803 has been safe in Phase I trials in humans, and is already in use in clinical trials in cancer patients (33, 34), as well as phase II clinical trials in HIV+ patients in Thailand and in the United States (35) (https://immunitybio.com/immunitybio-announces-launch-of-phase-2-trial-of-il-15-superagonist-anktiva-with-antiretroviral-therapy-to-inhibit-hiv-reservoirs/; https://actgnetwork.org/studies/a5386-n-803-with-or-without-bnabs-for-hiv-1-control-in-participants-living-with-hiv-1-on-suppressive-art/).

IL-15 can activate MAIT cells *in vitro* (19); however, no studies have been performed to evaluate the effect of IL-15 or N-803 on MAIT cells *in vivo*. Furthermore, the impact of IL-15 or N-803 treatment on the anti-mycobacterial function of MAIT cells is not known. We hypothesized that N-803 could activate MAIT cells *in vitro,* as well as improve MAIT cell activation, function, or trafficking to mucosal sites *in vivo*.

Macaques provide an ideal model system in which to characterize the effects of N-803 on MAIT cell trafficking and cytolytic function, as macaque and human MAIT cells are phenotypically and functionally similar (36, 37). To test our hypothesis, we first examined the *in vitro* effect of N-803 on MAIT cells isolated from healthy and Simian Immunodeficiency Virus (SIV)-positive macaques. Then, we examined MAIT cells from a previous study of SIV+, ART-naïve rhesus macaques who were treated with N-803. (26)(Ellis-Connell et al; MS in prep). We examined the effects of N-803 on MAIT cell phenotype and function in the PBMC, lymph nodes, airways, and lung tissue *in vitro* and *in vivo*.

## Materials and Methods

### Animals and reagents

#### Animals

The samples collected from SIV+, ART-naïve rhesus macaques treated *in vivo* with N-803 are described in detail in (Ellis et al., manuscript submitted). All animals involved in this study were cared for and housed at the Wisconsin National Primate Resource Center (WNPRC), following practices that were approved by the University of Wisconsin Graduate School Institutional Animals Care and Use Committee (IACUC; protocol number G005507). All procedures, such as bronchoalveolar lavage, biopsies, blood draw collection, and N-803 administration were performed as written in the IACUC protocol, under anesthesia to minimize suffering.

#### Frozen samples

Frozen PBMC from other SIV-naïve and SIV+ cynomolgus macaques were collected as parts of previous studies (14, 38, 39) and utilized for *in vitro* assays described here.

Table 1 shows the animals from which samples were utilized for this study, the SIV strain(s) they were challenged with, and which figures samples from that animal were used for.

**Table 1.** Animals and samples utilized in this study.

| Animal | SIV strain(s)<br>Challenged with | Study previously<br>described in | Figure(s) samples<br>were utilized in |
| --- | --- | --- | --- |
| Cy0899 | SIVmac239 | (14) | Fig 1 |
| Cy0900 | SIVmac239 | (14) | Figs 1, 2 |
| Cy0918 | SIVmac239 | (14) | Figs 1, 2 |
| Cy0919 | SIVmac239 | (14) | Figs 1, 2 |
| Cy0756 | SIVmac239Δnef-8X; SIVmac239 | (39) | Fig 2 |
| Cy0757 | SIVmac239Δnef-8X; SIVmac239 | (39) | Fig 2 |
| r15053 | SIVmac239M | Ellis-Connell et al,<br>2022; submitted | Figs. 3-8; 10 |
| r15090 | SIVmac239M | Ellis-Connell et al,<br>2022; submitted | Figs. 3-8; 10 |
| r03019 | SIVmac239M | Ellis-Connell et al,<br>2022; submitted | Figs. 3-8; 10 |
| rh2498 | SIVmac239M | Ellis-Connell et al,<br>2022; submitted | Figs. 3-8; 10 |
| rh2493 | SIVmac239M | Ellis-Connell et al,<br>2022; submitted | Figs. 3-8; 10 |
| rh2903 | SIVmac239M | Ellis-Connell et al,<br>2022; submitted | Figs. 4-10 |
| rh2906 | SIVmac239M | Ellis-Connell et al,<br>2022; submitted | Figs. 4-10 |
| rh2907 | SIVmac239M | Ellis-Connell et al, 2022; submitted | Figs. 4-10 |
| rh2909 | SIVmac239M | Ellis-Connell et al, 2022; submitted | Figs. 4-10 |
| rh2911 | SIVmac239M | Ellis-Connell et al, 2022; submitted | Figs. 4-10 |
| rh2920 | SIVmac239M | Ellis-Connell et al, 2022; submitted | Figs. 4-10 |
| rh2921 | SIVmac239M | Ellis-Connell et al, 2022; submitted | Figs. 4-10 |
| r05053 | SIVmac239M | Ellis-Connell et al, 2022; submitted | Figs. 4-10 |
| r10063 | SIVmac239M | Ellis-Connell et al, 2022; submitted | Figs. 4-10 |

#### N-803 reagent and administration

N-803 was provided by ImmunityBio (Culver City, CA), and produced using methods previously described (27, 30, 31). Briefly, all macaques treated with N-803 received three doses of 0.1mg/kg, delivered subcutaneously, separated by 14 days. For the present study, the effects of N-803 on MAIT cell immunology are only described for the first dose and 14 days following. This N-803 dose and route of administration were previously found to be safe and efficacious in macaques (26, 31)

### Sample collection for use in *ex vivo* MAIT cell characterization following N-803 administration

#### PBMC and Lymph node (LN)

Peripheral blood mononuclear cells (PBMC) from macaques from the *in vivo* study were isolated from whole blood by ficoll density gradient centrifugation as previously described (14, 26, 39). For all assays described in this manuscript, PBMC were frozen in CryoStor CS5 Freezing media (BioLife Solutions, Inc; Bothell, WA) and stained for phenotypic and functional characterization of MAIT cells. Lymph node biopsies were processed by cutting them into small pieces and manually passing them through a 70μm filter. Then cells were separated from the tissue by ficoll density gradient centrifugation as above. Cells were frozen and stained for phenotypic characterization of MAIT cells in bulk assays (described below).

#### Bronchoalveolar Lavage (BAL)

BAL fluid was collected by primate center staff as previously described (40). Briefly, 30mL of phosphate-buffered saline (PBS) solution was flushed into the airways of macaques and aspirated fully. BAL fluid was then passed through a 70μm filter. Cells were resuspended in RPMI media supplemented with 10% Fetal bovine serum (FBS) for use in *ex vivo* assays (described below).

#### Lung

Lung tissues were collected at the time of necropsy by primate center staff. Tissue samples were homogenized, then digested for 2 hours at 37 degrees celsius in a solution consisting of RPMI supplemented with 10% FBS and 0.1mg/mL collagenase II (Sigma Aldrich; St. Louis, MO). After digestion, homogenates were passed through a 70μm filter and resuspended in fresh media containing 35% isotonic percoll. Cells were then layered over 60% isotonic percoll and were isolated by percoll gradient centrifugation (1860 RCF for 30 minutes). Lymphocytes were collected from the interphase between the 35% and 60% layers, resuspended in RPMI media supplemented with 10% FBS, and used in *in vitro* assays described below.

### Flow cytometric analysis

#### MR1-5-OP-RU Tetramer reagents

The rhesus macaque MR1-5OPRU monomer was provided by the NIH tetramer core facility. The MR1 tetramer technology was developed jointly by Dr. James McCluskey, Dr. Jamie Rossjohn, and Dr. David Fairlie, and the material was produced by the NIH Tetramer Core Facility as permitted to be distributed by the University of Melbourne. The MR1-5-OP-RU monomer was then tetramerized with streptavidin-BV421 (0.1mg/mL; BD biosciences) or streptavidin-APC (0.74mg/mL; Agilent technologies) at an 8:1 molar ratio of monomer:streptavidin. Briefly, 1/10th volumes of streptavidin-BV421 or streptavidin-APC were added to the monomer every 10 minutes and incubated in the dark at 4°C until the 8:1 molar ratio was achieved.

#### Staining methods

For all flow cytometry panels (antibodies described in Tables 2-7), the order of staining was as follows: surface staining with the MR1-5-OP-RU tetramer was performed prior to other surface and intracellular staining. MR1 tetramer stains were performed at room temperature in the dark for 45 minutes in RPMI media supplemented with 10% FBS and 50nM dasatinib (Thermo Fisher Scientific, Waltham, MA). After 45 minutes, the cells were washed in a solution of FACS buffer (2% FBS in a 1X PBS solution) containing 50nM dasatinib, and surface stains were performed using the antibodies indicated in Tables 2-7 for 20 minutes at room temperature in the dark. Cells were fixed in a 2% paraformaldehyde solution. Following a 20 minute incubation, samples were either run on a BD Symphony A3 or permeabilized and stained for 20 minutes at room temperature in medium B (Thermo Fisher Scientific, Waltham, MA) for intracellular markers. Flow cytometric analysis was performed on a BD Symphony A3 (Becton Dickinson, Franklin Lakes, NJ), and the data were analyzed using FlowJo software for Macintosh (version 10.7.1).

**Table 2.** Cytolytic granule/proliferation flow cytometry panel used for PBMC

| Antibody | Clone | Tetramer/Surface/Intracellular |
| --- | --- | --- |
| Gag<sub>181-189</sub>CM9-PE | --- | Tetramer*** |
| Tat<sub>28-35</sub>SL8-BV605 | --- | Tetramer*** |
| MR1 5-OP-RU-BV421 | --- | Tetramer |
| CD3-AF700 | SP34-2 | Surface |
| CD4-BUV737 | SK3 | Surface |
| CD8-BUV395 | RPA-T8 | Surface |
| CD28-BV510 | CD28.2 | Surface |
| CD95-PE Cy7 | DX2 | Surface |
| CCR7-BV711 | G043H7 | Surface |
| LIVE/DEAD Near IR ARD | --- | --- |
| ki-67-AF488 | B56 | Intracellular |
| GranzymeB-PE CF594 | GB11 | Intracellular |
| Perforin-AF647 | PF344 | Intracellular |

\*\*\*The Gag<sub>181-189</sub>CM9-PE and Tat<sub>28-35</sub>SL8-BV605 tetramers were used to characterize SIV-specific conventional T cells, which are described elsewhere (manuscript in progress).

**Table 3.** Chemokine/trafficking flow cytometry panel used for PBMC

| Antibody | Clone | Tetramer/Surface |
| --- | --- | --- |
| Gag<sub>181-189</sub>CM9-PE | --- | Tetramer*** |
| Tat<sub>28-35</sub>SL8-BV605 | --- | Tetramer*** |
| MR1 5-OP-RU-APC | --- | Tetramer |
| CD3-AF700 | SP34-2 | Surface |
| CD4-BUV737 | SK3 | Surface |
| CD8-BUV395 | RPA-T8 | Surface |
| CXCR3-BV650 | G025H7 | Surface |
| CCR6-PE Cy7 | 11A9 | Surface |
| CXCR5-PerCP Efluor710 | MU5UBEE | Surface |
| CD122-BV421 | Mik-β3 | Surface |
| CD132-BB515 | AG184 | Surface |
| LIVE/DEAD Near IR ARD | --- | --- |

\*\*\*The Gag<sub>181-189</sub>CM9-PE and Tat<sub>28-35</sub>SL8-BV605 tetramers were used to characterize SIV-specific conventional T cells, which are described elsewhere (manuscript in progress).

**Table 4.**
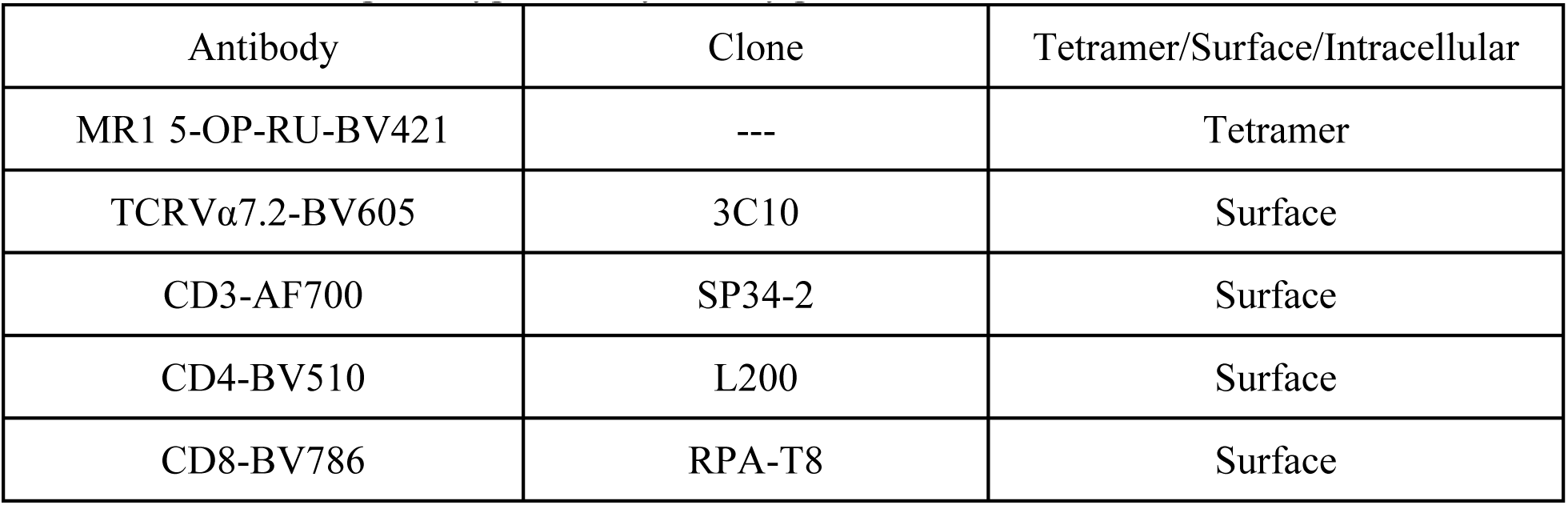
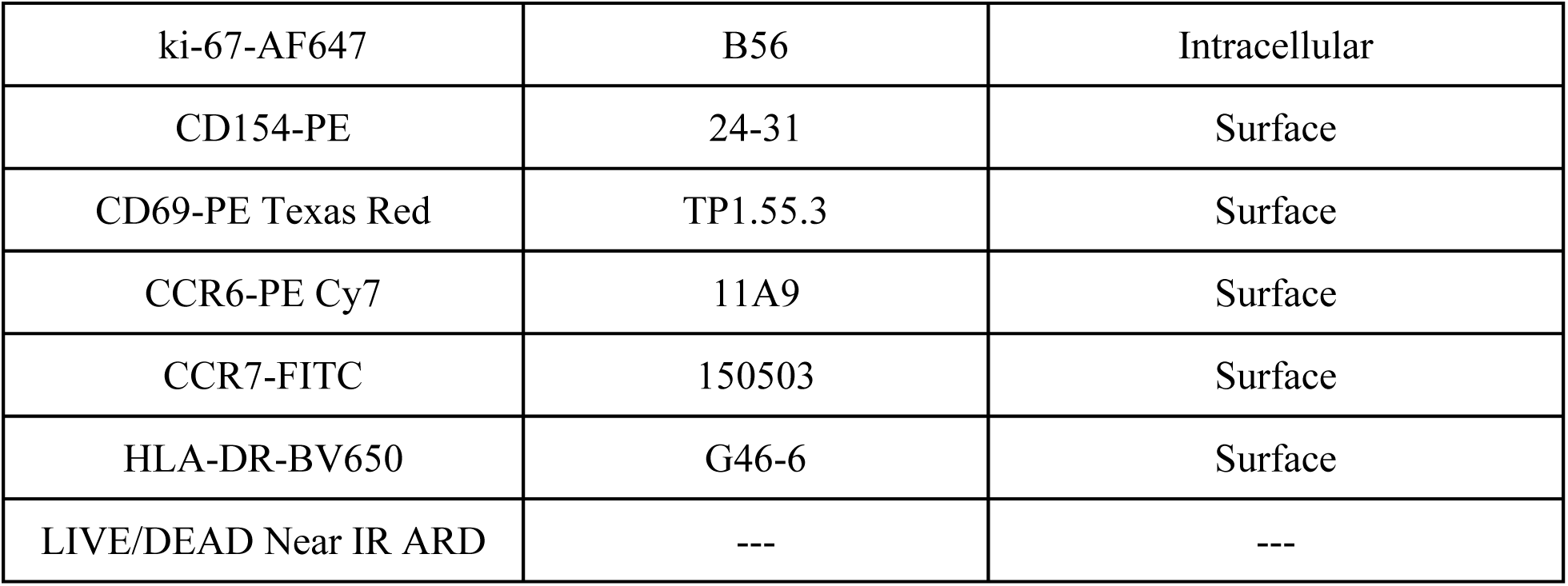
MAIT BAL phenotype flow cytometry panel

### *In vitro* MAIT cell activation assays with N-803

#### MAIT cell isolation

MAIT cells were isolated from cryopreserved PBMC of SIV-naïve macaques by adding 10μL of TCRVα7.2-PE (clone 3C10, Biolegend; San Diego, CA) for every 5e6 cells, and incubating for 20 minutes at 4°C. Then, PE-labeled cells were isolated using a MACS Miltenyi PE microbeads kit according to manufacturer’s protocols (130-097-054; MACS Miltenyi, Auburn CA). Following isolation, cells were used for *in vitro* N-803 activation assays (described below).

#### N-803 administration and staining for activation markers on MAIT cells

Bulk cryopreserved PBMC from SIV-naïve macaques or MAIT cells isolated using TCRVα7.2-PE (described above) from frozen PBMC were suspended in RPMI supplemented with 10% fetal bovine serum (FBS) and 100units/mL penicillin/streptomycin (Thermo Fisher Scientific; Waltham, MA); and 2mM L-glutamine (Thermo Fisher Scientific; Waltham, MA). The indicated concentrations of N-803, or recombinant human IL-15 (rhIL15, Peprotech; Cranbury, NJ), were added to the media, then incubated overnight at 37°C. The following day, the cells were collected and stained with the antibodies indicated in the panel below (Table 5). Flow cytometry was performed as described above.

**Table 5.** Antibodies used for *in vitro* MAIT activation assay

| Antibody | Clone | Tetramer/Surface |
| --- | --- | --- |
| MR1 5-OP-RU-BV421 | ----- | Tetramer |
| CD8-BUV563 | RPA-T8 | Surface |
| CD4-BUV737 | SK3 | Surface |
| CD3-BUV395 | SP34-2 | Surface |
| CD69-PE Texas Red | TP1.55.3 | Surface |
| HLA-DR BV650 | G46-6 | Surface |
| CD25-APC | BC96 | Surface |
| LIVE/DEAD Near IR ARD | ----- | ----- |

### *In vitro* and *ex vivo* MAIT cell functional assays

#### In vitro MAIT cell functional assays

Functional assays were performed using cryopreserved PBMC and LN from SIV-naïve and SIV+ macaques, and freshly isolated lymphocytes from the lungs of SIV+ macaques. Cells were incubated overnight at 37°C in 96-well plates in RPMI supplemented with 15% FBS, along with the indicated concentrations of N-803, recombinant IL-7 (rhIL-7; Biolegend; San Diego, CA), or media control. The following day, the PBMC that were incubated overnight in various cytokines or media control were counted, and functional assays were performed. Briefly, fixed *E.coli* and *M.smegmatis* stimuli were prepared as described in (14, 41), by fixing bacteria in 2% paraformaldehyde for 2-3 minutes, followed by three washes with 1X phosphate buffered saline (PBS), then resuspension in RPMI media supplemented with 15% FBS. Ten colony-forming units (CFU) of fixed *E.coli* or *M.smegmatis* were added for every one cell present per well. The cells were incubated with the bacteria for 90 minutes at 37°C, then brefeldin A and monensin were added along with 5uL of CD107a-BV605. The cells were incubated for another 6 hours, then staining was performed as indicated above, using the antibodies listed in Table 6. Flow cytometry was performed as described above.

**Table 6.** MAIT *in vitro* and *ex vivo* functional assays.

| Antibody | Clone | Tetramer/Surface/Intracellular |
| --- | --- | --- |
| MR1 5-OP-RU-BV421 | ----- | Tetramer |
| CD3-BUV396 | SP34-2 | Surface |
| CD4-BUV737 | SK3 | Surface |
| CD8-BUV563 | RPA-T8 | Surface |
| CD107a-BV605 | H4A3 | Surface |
| IFNγ-FITC | 4S.B3 | Intracellular |
| TNFα-AF700 | Mab11 | Intracellular |
| LIVE/DEAD Near IR ARD | ----- | ----- |

**Table 7.**
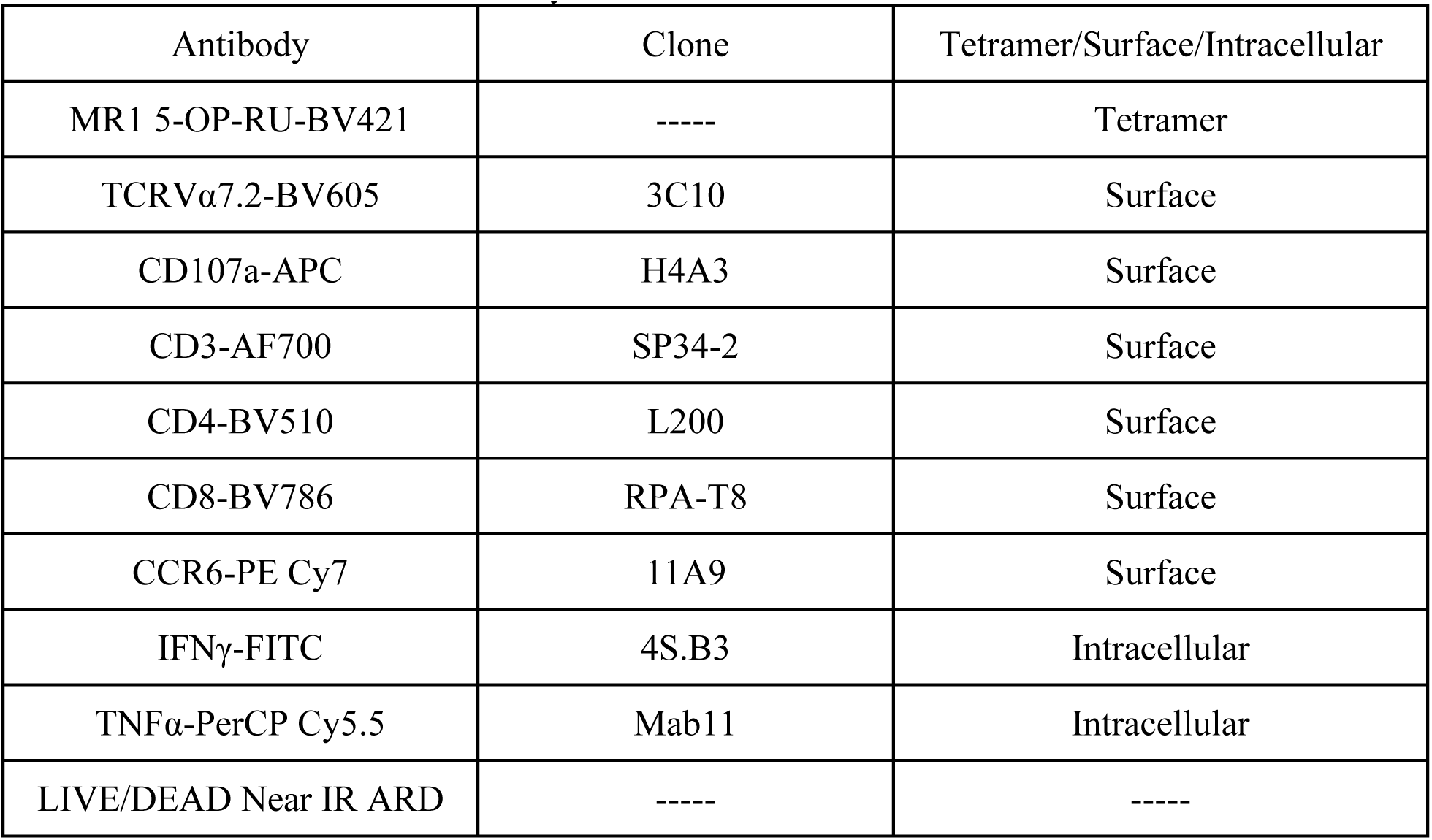
MAIT BAL functional assay

| Antibody | Clone | Tetramer/Surface/Intracellular |
| --- | --- | --- |
| MR1 5-OP-RU-BV421 | ----- | Tetramer |
| TCRVα7.2-BV605 | 3C10 | Surface |
| CD107a-APC | H4A3 | Surface |
| CD3-AF700 | SP34-2 | Surface |
| CD4-BV510 | L200 | Surface |
| CD8-BV786 | RPA-T8 | Surface |
| CCR6-PE Cy7 | 11A9 | Surface |
| IFNγ-FITC | 4S.B3 | Intracellular |
| TNFα-PerCP Cy5.5 | Mab11 | Intracellular |
| LIVE/DEAD Near IR ARD | ----- | ----- |

#### Ex vivo MAIT cell functional assays performed with cryopreserved PBMC collected from N-803 treated macaques

Cryopreserved PBMC originally collected from macaques treated with N-803 were thawed from the indicated time points pre- and post-N-803 treatment. Thawed cells were rested for 4 hours. Then, 10 CFU of fixed *E.coli* or *M.smegmatis* were added for every one cell present in the tube. Cells and bacteria were incubated together for 90 minutes at 37°C, and then brefeldin A, monensin, and CD107a-BV605 were added. Cells were incubated overnight at 37°C. The following day, cells were stained as indicated in Table 6, and flow cytometric analysis was performed as described above.

#### Ex vivo MAIT cell functional assays performed with freshly isolated BAL collected from N-803 treated macaques

BAL fluid was collected and processed as indicated above. Then, 10 CFU of either fixed *E.coli* or *M.smegmatis* were added for every one cell present to the appropriate tubes as described above. Cells and bacteria were incubated for 90 minutes at 37°C, then brefeldin A, monensin, and CD107a-APC were added. Cells were incubated for 6 hours at 37°C, then stained as indicated in Table 7. Flow cytometry was performed as described above.

## Statistical analysis

For statistical analyses performed with the same individuals across multiple timepoints, repeated measures ANOVA non-parametric tests were performed, with Dunnett’s multiple comparisons (Prism GraphPad). For individuals with missing timepoints, mixed-effects ANOVA tests were performed using Geisser-Greenhouse correction.

For statistical analyses in which two timepoints were being compared to each other for the same individuals, Wilcoxon matched pairs signed rank tests were performed.

## Results

### N-803 can activate MAIT cells from macaque PBMC

We first wanted to determine if N-803 could activate MAIT cells, *in vitro*, at similar or greater levels when compared to rhIL-15. RhIL-15 activates MAIT cells directly through the IL-15 receptor, and indirectly through the production of IL-18 by monocytes, which activates MAIT cells via the IL-18 receptor (19). Therefore, to assess both the direct and indirect impact of N-803 on MAIT cell activation, we used total PBMC from healthy cynomolgus macaques or isolated the TCRVα7.2+ cells (MAIT cells in these macaques are primarily TCRVα7.2+ (14)). Then, total PBMC or TCRVα7.2+ cells were treated for 24 hours with increasing concentrations of recombinant rhIL-15 or N-803 (Fig. 1A). We then measured the frequency of MAIT cells expressing the activation markers CD69, CD25, and HLA-DR (Fig. 1B shows an example gating schematic).

**Fig. 1.**
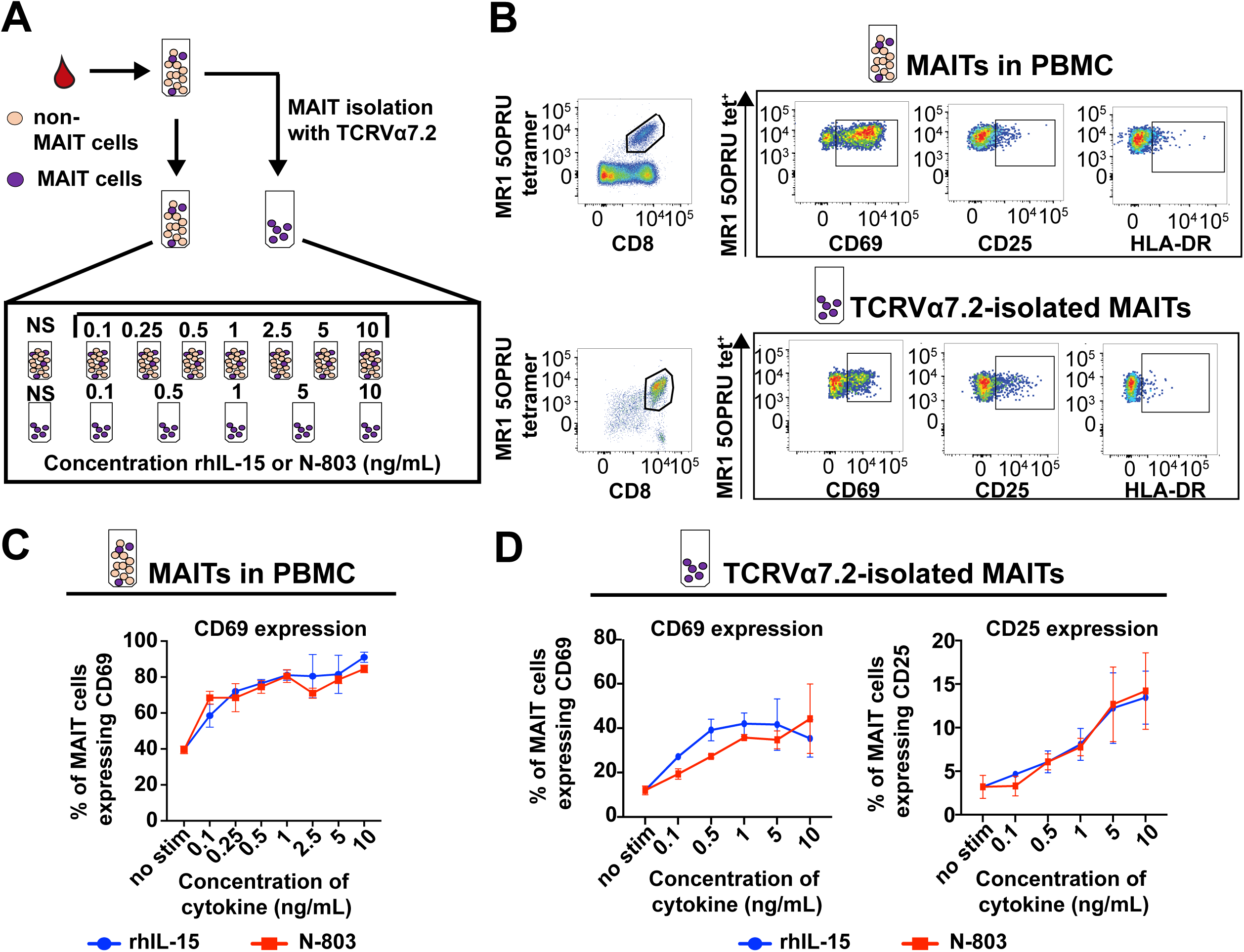
IL-15 superagonist N-803 activates MAIT cells *in vitro*. A, Schematic of *in vitro* experiments performed. Frozen PBMC from healthy macaques were incubated overnight with the indicated concentrations of either recombinant IL-15 (rhIL-15) or N-803. Alternatively, the TCRVα7.2+ cells (MAIT cells, purple dots) were isolated from PBMC, and then incubated overnight with the indicated concentrations of either rhIL-15 or N-803. Flow cytometry was performed after incubation to assess activation. B, Gating schematic used for flow cytometry for the experiments described in (A) to detect MAIT cells present in PBMC (top panels) or TCRVα7.2-isolated cells (bottom panels), and the frequencies of CD69+, CD25+, and HLA-DR+ cells. C, Graphical analysis of the experiments described in (A) for MAIT cells present in PBMC. The frequencies of MAIT cells expressing CD69 are shown for the indicated concentrations of rhIL-15 (blue circles) or N-803 (red squares). D, Graphical analysis of the experiments described in (A) for TCRVα7.2-isolated MAIT cells. Graphs show the frequencies of MAIT cells expressing either CD69 (left) or CD25 (right) for the indicated concentrations of rhIL-15 (blue circles) or N-803 (red squares).

We found that the addition of rhIL-15 or N-803 to PBMC led to a dose-dependent increase in the frequency of CD69+ MAIT cells (Fig. 1C). We did not observe any increases in the frequency of MAIT cells expressing CD25 or HLA-DR (data not shown). When using isolated TCRVα7.2+ cells, we found dose-dependent increases in the frequency of CD69+ (Fig. 1D, left graph) and CD25+ (Fig. 1D, right graph) cells. These data suggest that like rhIL-15, N-803 has direct and indirect impacts on activation of MAIT cells *in vitro*.

### N-803 improves IFNγ production of bacterial-stimulated MAIT cells derived from the PBMC of SIV+ and SIV-naïve cynomolgus macaques

We wanted to determine if N-803 enhanced the production of IFNγ, TNFα, and CD107a by MAIT cells stimulated *in vitro* with *E. coli* or *M. smegmatis*. We included PBMC from cynomolgus macaques collected before infection with various strains of SIV, and then PBMC collected from the same macaques at necropsy, after SIV infection (Table 1). PBMC were incubated overnight in media alone, N-803, or recombinant IL-7. Recombinant IL-7 has been shown increase the frequency of cytolytic MAIT cells when co-incubated with antigen (21). The following day, 10 colony forming units (CFU) of either *E.coli* or *M.smegmatis* were then added for every one cell and incubated together for another six hours. We measured the frequencies of IFNγ, TNFα, and CD107a+ MAIT cells by flow cytometry. An example of the gating schematic and typical cytokine and cytolytic enzyme production during bacterial stimulation can be found in Supplementary Fig. 1.

We found that incubation with N-803 enhanced the frequency of MAIT cells producing IFNγ, TNFα, and CD107a after stimulation with *E.coli* in samples from SIV-naïve timepoints. Only the frequencies of MAIT cells producing IFNγ+ and TNFα+ were improved with N-803 treatment after stimulation with *M.smegmatis* (Fig. 2A). For PBMC collected after SIV infection, N-803 treatment enhanced IFNγ production from MAIT cells for both *E.coli* and *M.smegmatis* stimuli but did not significantly enhance TNFα or CD107a production in a consistent manner (Fig. 2B). Together, these data suggest that N-803 treatment improves the cytokine production of MAIT cells incubated with microbial stimuli, albeit to a lesser extent in PBMC collected from SIV+ macaques.

**Fig. 2.**
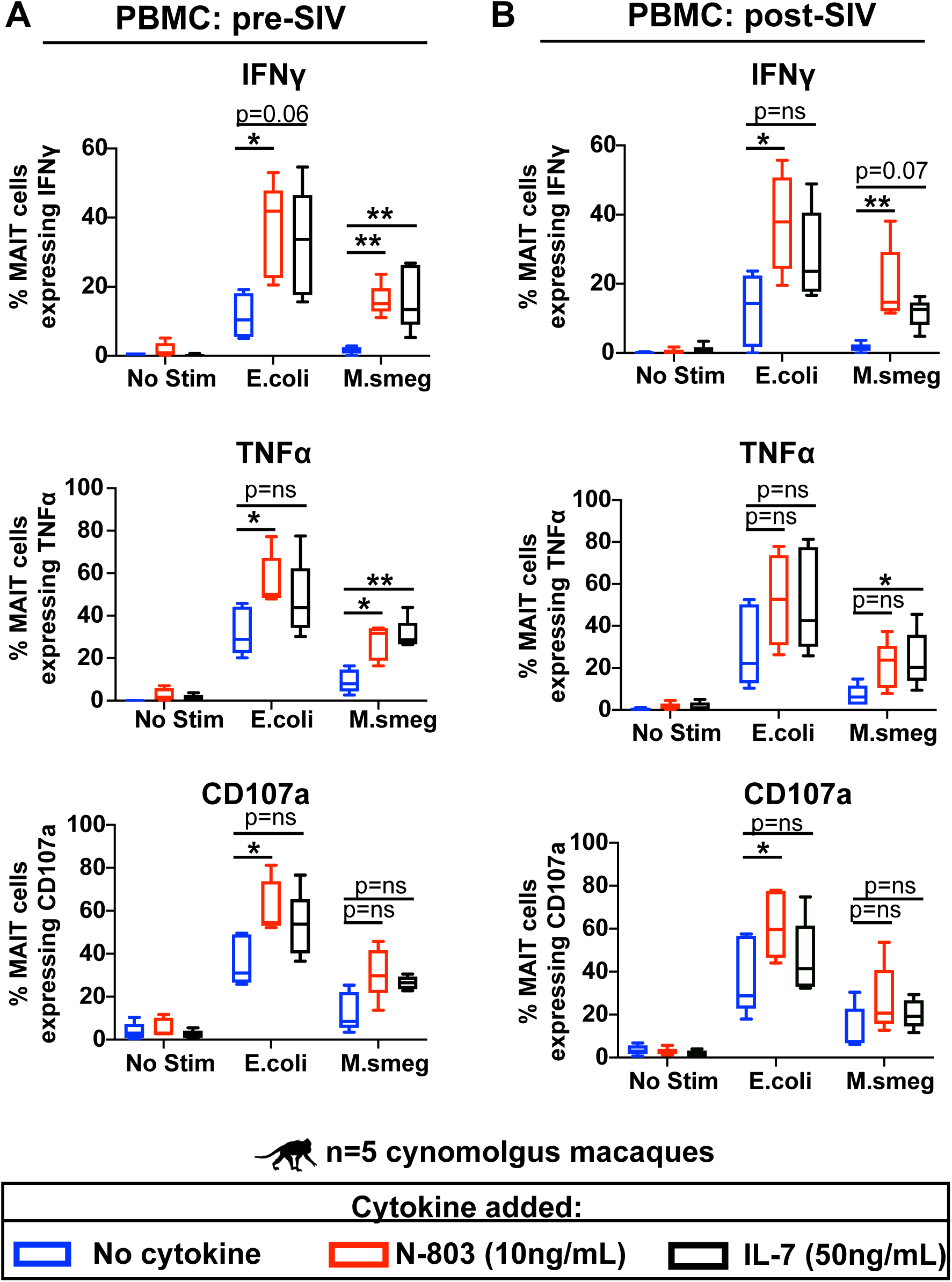
N-803 increases IFNγ production from MAIT cells stimulated with either *E.coli* and *M.smegmatis* from both SIV-naive and SIVmac239+ macaques. A, Frozen PBMC from healthy cynomolgus macaques (n=5) were incubated overnight with 10ng/mL N-803, 50ng/mL IL-7, or media control. The following day, functional assays were performed using 10 CFU bacteria/cell of either *E.coli* or *M.smegmatis* as stimuli as described in the methods. Cells were incubated for 6 hours in the presence of bacteria and then flow cytometry was performed to determine the frequencies of MAIT cells expressing IFNγ (top), TNFα (middle), or CD107a (bottom). ANOVA tests with Dunnett’s multiple comparisons were performed to determine statistical significance between each treatment group (no stim, N-803, or IL-7) for each stimulus (*E.coli* or *M.smegmatis*). p=ns; not significant, *; p≤0.05; **; p≤0.005. B, Frozen PBMC from the same cynomolgus macaques (n=5) as in (A) were taken from necropsy timepoints, which occurred 6 months to 1 year after SIVmac239 infection. PBMC were incubated with 10ng/mL N-803, 50ng/mL IL-7, or media control as in (A). Functional assays were performed as described in (A) to determine the frequencies of MAIT cells expressing IFNγ (top), TNFα (middle), or CD107a (bottom). ANOVA tests with Dunnett’s multiple comparisons were performed to determine statistical significance between each treatment group (no stim, N-803, or IL-7) for each stimulus (*E.coli* or *M.smegmatis*). p=ns; not significant, *; p≤0.05; **; p≤0.005.

### N-803 improves IFNγ production of bacterial-stimulated MAIT cells derived from the PBMC and lung tissue, but not lymph nodes, of SIV+ rhesus macaques

MAIT cells located in lung tissue and lung-associated lymph nodes may not behave the same as those derived from the PBMC (42, 43). We isolated lymphocytes from blood (PBMC), lung tissue, and tracheobronchial lymph nodes collected at necropsy from 4 SIV-infected rhesus macaques (Ellis-Connell et al, manuscript in progress; Fig. 3). Assays with lung lymphocytes were performed using fresh samples, while cryopreserved PBMC and lymph node cells were used.

**Fig. 3.**
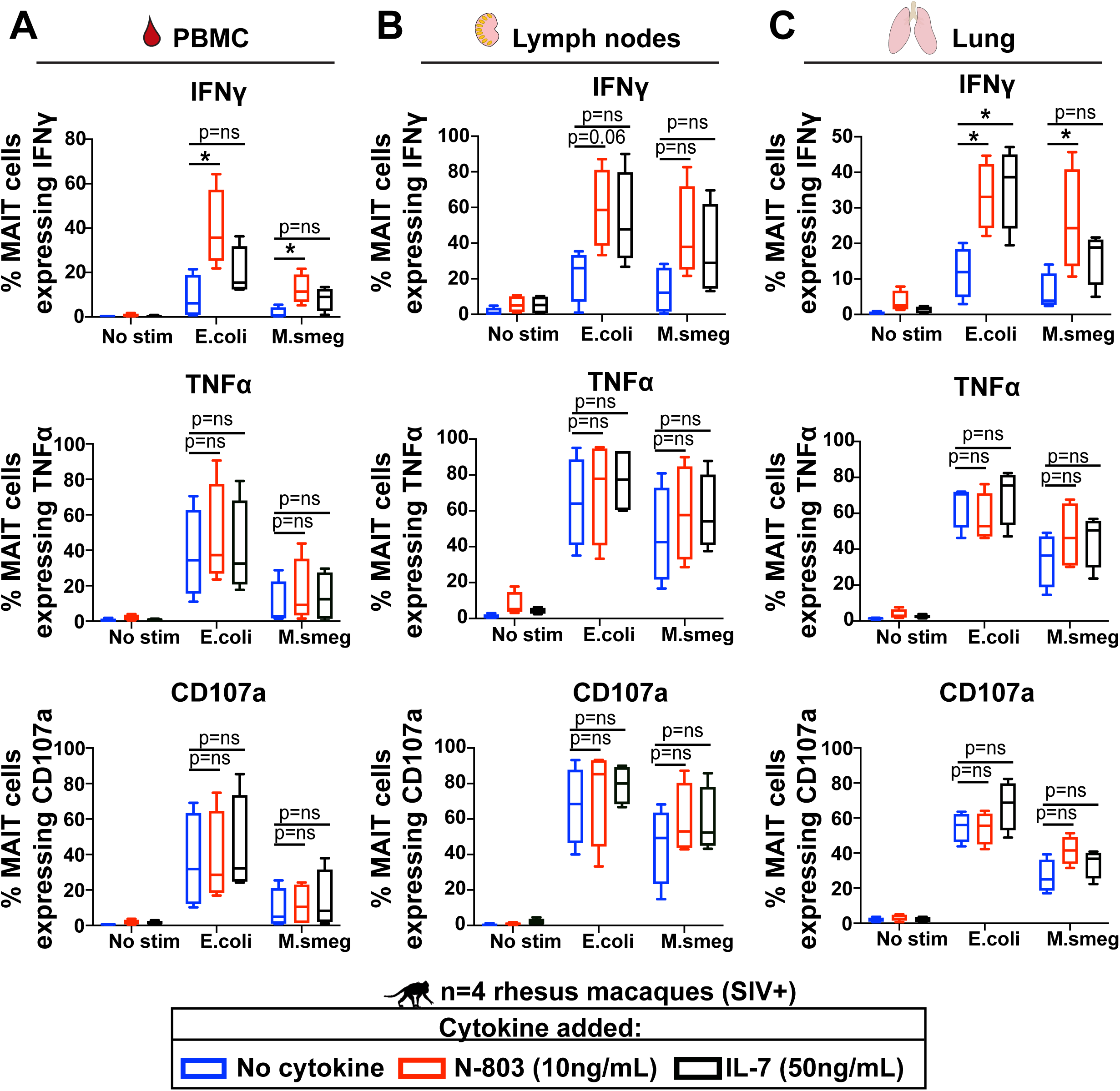
*In vitro* treatment of MAIT cells present in PBMC and lung tissue, but not lymph nodes, with N-803 increases IFNγ production with *E.coli* and *M.smegmatis* bacterial stimuli. SIVmac239+ rhesus macaques (n=4) were necropsied after greater than 8 months of infection, and lymphocytes were isolated from blood, tracheobronchial lymph nodes, and lung tissue. Frozen lymphocytes from PBMC (A), lymph nodes (B), and fresh lymphocytes from lung (C) were incubated overnight with 10ng/mL N-803, 50ng/mL IL-7, or media control. The following day, functional assays were performed using 10 CFU bacteria/cell of either *E.coli* or *M.smegmatis* as stimuli as described in the methods. Flow cytometry was performed to determine the frequencies of MAIT cells expressing IFNγ (top), TNFα (middle), or CD107a (bottom). ANOVA tests with Dunnett’s multiple comparisons were performed to determine statistical significance between each treatment group (no stim, N-803, or IL-7) for each stimulus (*E.coli* or *M.smegmatis*). p=ns; not significant, *; p≤0.05.

Similar to the data from the SIV+ cynomolgus macaques (Fig. 2B), incubation of PBMC from SIV+ rhesus macaque with N-803 led to statistically significant increases in the frequency of MAIT cells producing IFNγ with both *E.coli* and *M.smegmatis* stimuli when compared to media control (Fig. 3A). In the tissues, N-803 enhanced the frequency of IFNγ -producing bacterial-stimulated MAIT cells from the lung, but not the thoracic lymph nodes (Fig. 3B & C). In contrast, the frequency of MAIT cells producing TNFα or CD107a after microbial stimulation was unaffected by N-803 or IL-7 treatment in any compartment.

### Administration of N-803 to SIV+ rhesus macaques transiently decreases the number of peripheral MAIT cells

We wanted to determine how the administration of N-803 to macaques affected the frequency, distribution, and function of MAIT cells *in vivo*. We administered N-803 subcutaneously to ART-naïve, SIV+ rhesus macaques in a previous study (Fig. 4; Ellis et al., manuscript in preparation). We assessed the phenotypes of MAIT cells from the blood, bronchoalveolar lavage (BAL) fluid, and lymph nodes collected before and after receiving N-803 (Fig. 4). Due to the COVID-19 pandemic, the pre N-803 timepoints were collected between 14 and 42 days prior to N-803 treatment.

**Fig. 4.**
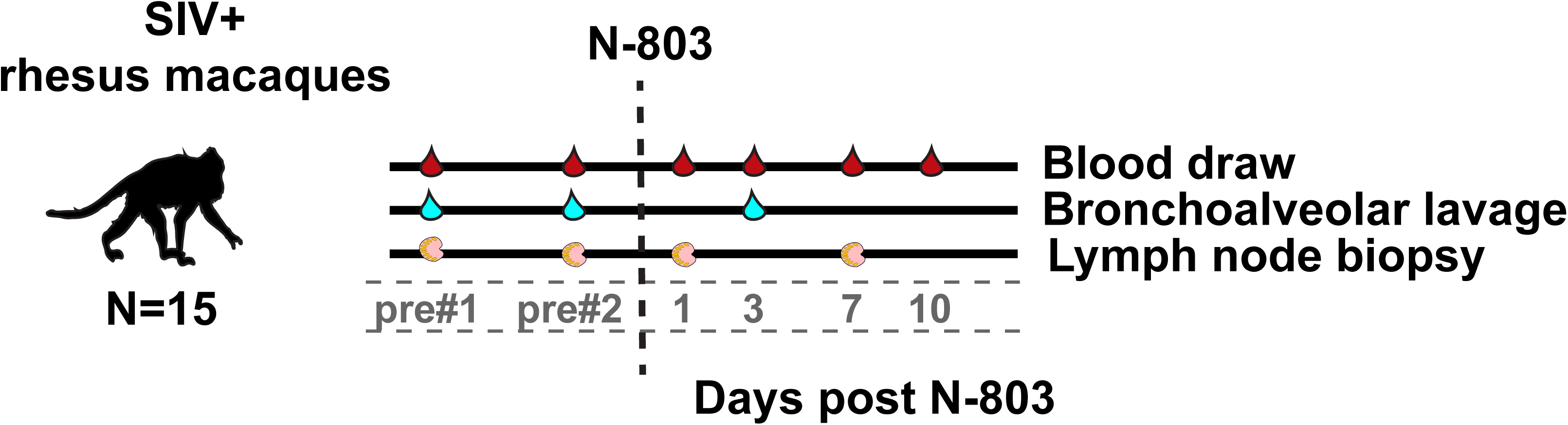
Study outline for N-803 treatment and sample collection. SIV+ rhesus macaques were given 0.1mg/kg N-803 subcutaneously. For the present study, samples that were collected prior to and in the first 10 days after the first dose of N-803 were used for analysis.

We defined MAIT cells as CD8+ MR1 tetramer+ cells (Fig. 5A). We found the frequencies of CD8+MR1 tet+ cells significantly decreased one day after N-803 treatment, but otherwise remained relatively stable relative to the average frequency prior to N-803 treatment (Fig 5B). The absolute number of MAIT cells in the blood were significantly lower on both days one and three post N-803 compared to the pre N-803 timepoints (Fig. 5C). We normalized the data relative to the average values of the pre N-803 timepoints on a per-animal basis to calculate the fold-change in frequency and absolute cell number (Figs. 5D and E). After normalization, we observed similar decreases on days one and three post N-803 (Figs. 5D and E). MAIT cells expanded in the blood on days 7 and 10 post N-803 treatment, but this was not statistically significant (Figs. 5D and E).

**Fig. 5.**
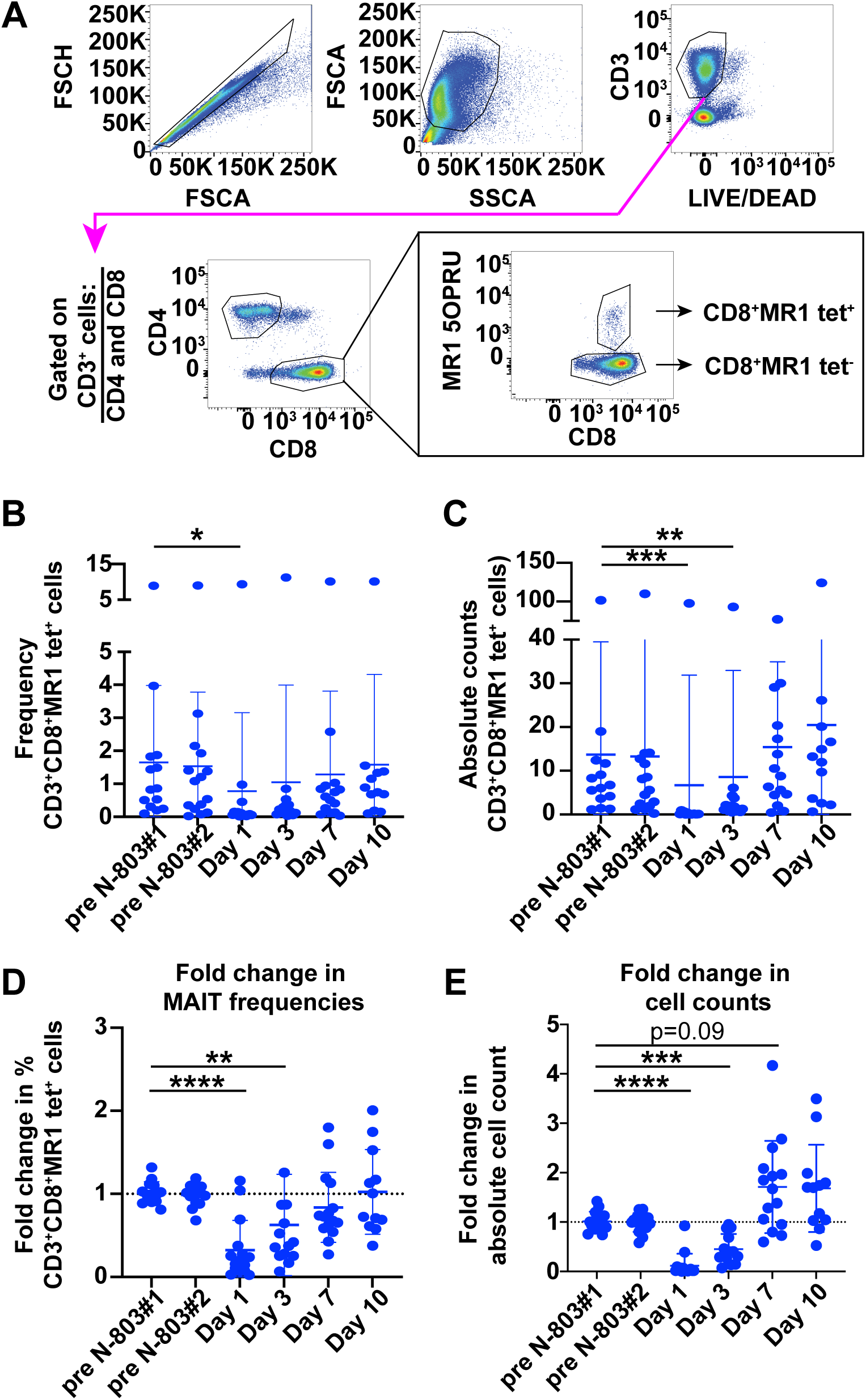
*In vivo* treatment of SIV+ macaques with N-803 alters MAIT cell frequencies in peripheral blood. A, PBMC collected from the indicated timepoints pre and post N-803 treatment were stained with the panel described in Table 3. Flow cytometry was performed as described in the methods. Shown is a representative gating schematic for CD3+CD8+MR1 tetramer+ (MAIT) cells. B, Flow cytometry was performed as described in (A) at the indicated time points post N-803 to determine the frequency of MAIT cells. C, Complete white blood cell counts (CBC) were used to quantify the absolute number of MAIT cells per μL of blood. D, The frequencies of MAIT cells from (B) were normalized to the pre N-803 averages for each animal, and the data are presented as fold change in the frequency of MAIT cells. E, The absolute counts of MAIT cells from (C) were normalized relative to the absolute cell counts of the pre N-803 averages for each animal and the data are presented as fold change in cell counts. For all statistical analysis in A-E, repeated measures ANOVA non-parametric tests were performed, with Dunnett’s multiple comparisons for individuals across multiple timepoints. For individuals for which samples from timepoints were missing, mixed-effects ANOVA tests were performed using Geisser-Greenhouse correction. *, p≤0.05; **, p≤0.005; ***, p≤0.0005; ****, p≤0.0001.

### N-803 treatment increases the frequency of Granzyme B and ki-67+ MAIT memory cells in the peripheral blood

Similar to the effects of N-803 on CD8 T cells and NK cells (23, 26, 31, 32), we expected that treatment of macaques with N-803 could potentially improve MAIT cell function. To begin to assess this, we measured the frequencies of MAIT cells expressing the degranulation marker Granzyme B and the proliferation marker ki-67. We separately examined the frequencies of MAIT cells expressing markers consistent with central memory (CM; CD28+CD95+) and effector memory (EM; CD28-CD95+) because recombinant IL-15 can differentially affect these two memory populations (44–49).

Similar to our previous study in cynomolgus macaques (14), peripheral MAIT cells from these rhesus macaques had both central and effector memory phenotypes (Fig 6A and B). The frequencies of CM and EM MAITs did not change during N-803 treatment (Fig. 6A and B, left). We next assessed the frequencies of CM and EM MAIT cells that were ki-67+ and Granzyme B+ over the course of N-803 treatment. We found transient increases in the frequencies of ki-67+ (Fig. 6A and B, middle) and Granzyme B+ (Fig. 6A and B, right) CM and EM MAIT cells on day 3 post N-803 treatment compared to pre N-803 timepoints. Overall, we concluded that N-803 treatment led to an increase in the frequency of MAIT cells capable of proliferating and degranulating.

**Fig. 6.**
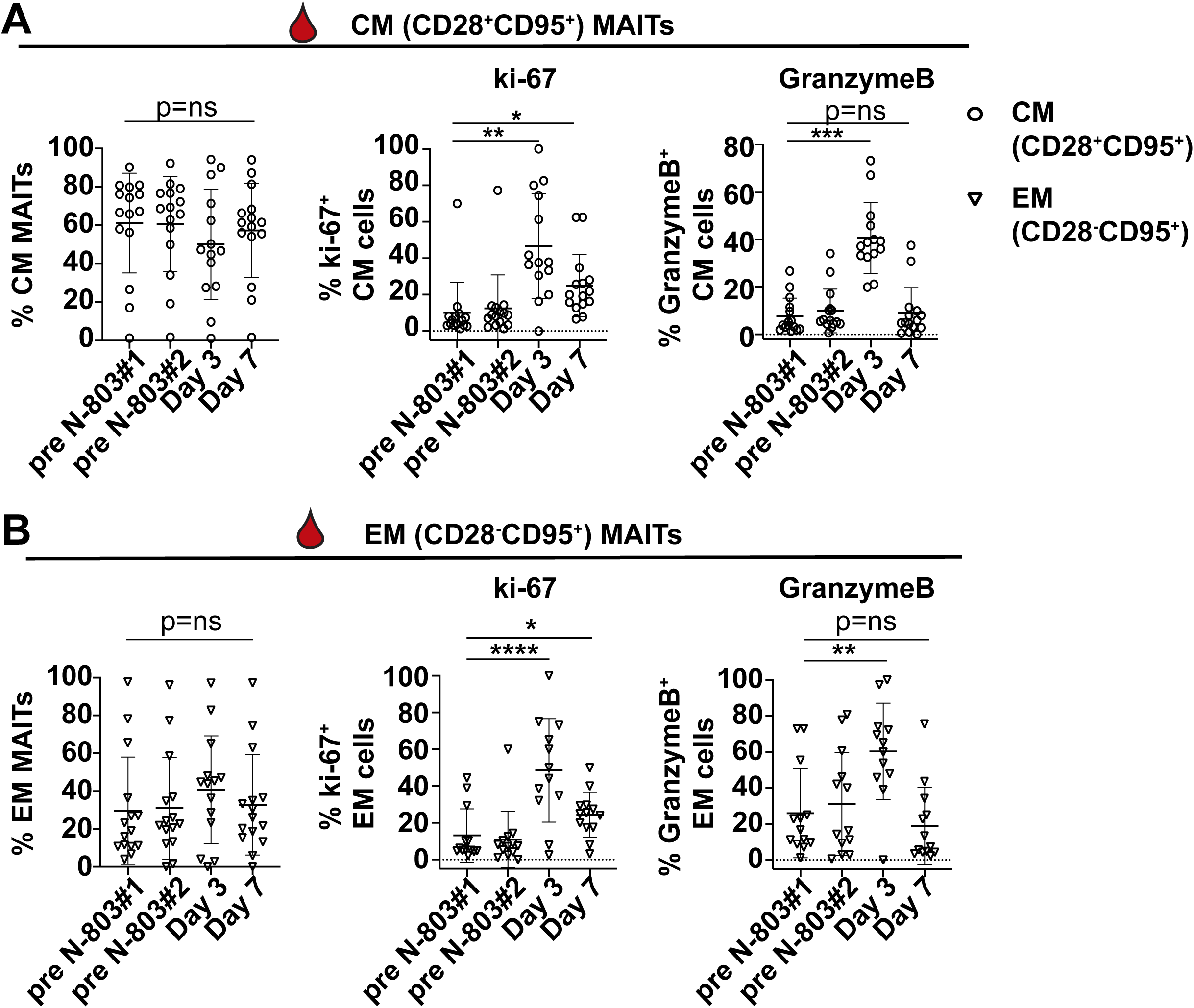
*In vivo* treatment of SIV+ macaques with N-803 increases the frequency of ki-67+ and Granzyme B+ MAIT cells in the peripheral blood. A and B, Frozen PBMC collected from the indicated timepoints pre and post N-803 were thawed, and flow cytometry was performed with the panel described in Table 2. The frequency of MAIT cells (CD8+MR1 tet+ cells) that were (A) central memory (CM; CD28+CD95+, open circles, left panel) or (B) effector memory (EM; CD28-CD95+, open triangles, left panel) are shown. The frequencies of CM and EM cells expressing ki-67 (A and B, middle panels) and Granzyme B (A and B, right panels) cells were determined. Repeated measures ANOVA non-parametric tests were performed, with Dunnett’s multiple comparisons for individuals across multiple timepoints. For individuals for which samples from timepoints were missing, mixed-effects ANOVA tests were performed using Geisser-Greenhouse correction. *, p≤0.05; **, p≤0.005.

### *In vivo* N-803 treatment leads to a decline in the frequency of peripheral CXCR3+CCR6+ MAIT cells

We hypothesized the decline in MAIT cells in the peripheral blood (Fig. 5) could be a consequence of MAIT cells trafficking to lymph nodes. We measured the frequency of CXCR5+ MAIT cells, as CXCR5 expression is associated with homing to lymph node follicles, which is similar to the traffic pattern of bulk CD3+CD8+ T cells observed in previous N-803 studies (23, 26, 31, 50–52). We also measured the frequency of MAIT cells expressing CXCR3 and CCR6, which have been associated with homing to tissues such as lung, gut, or other sites of immune activation (53–55). A representative gating schematic for the expression of these markers on MAIT cells prior to and 3 days post N-803 treatment is shown (Fig 7A).

**Fig. 7.**
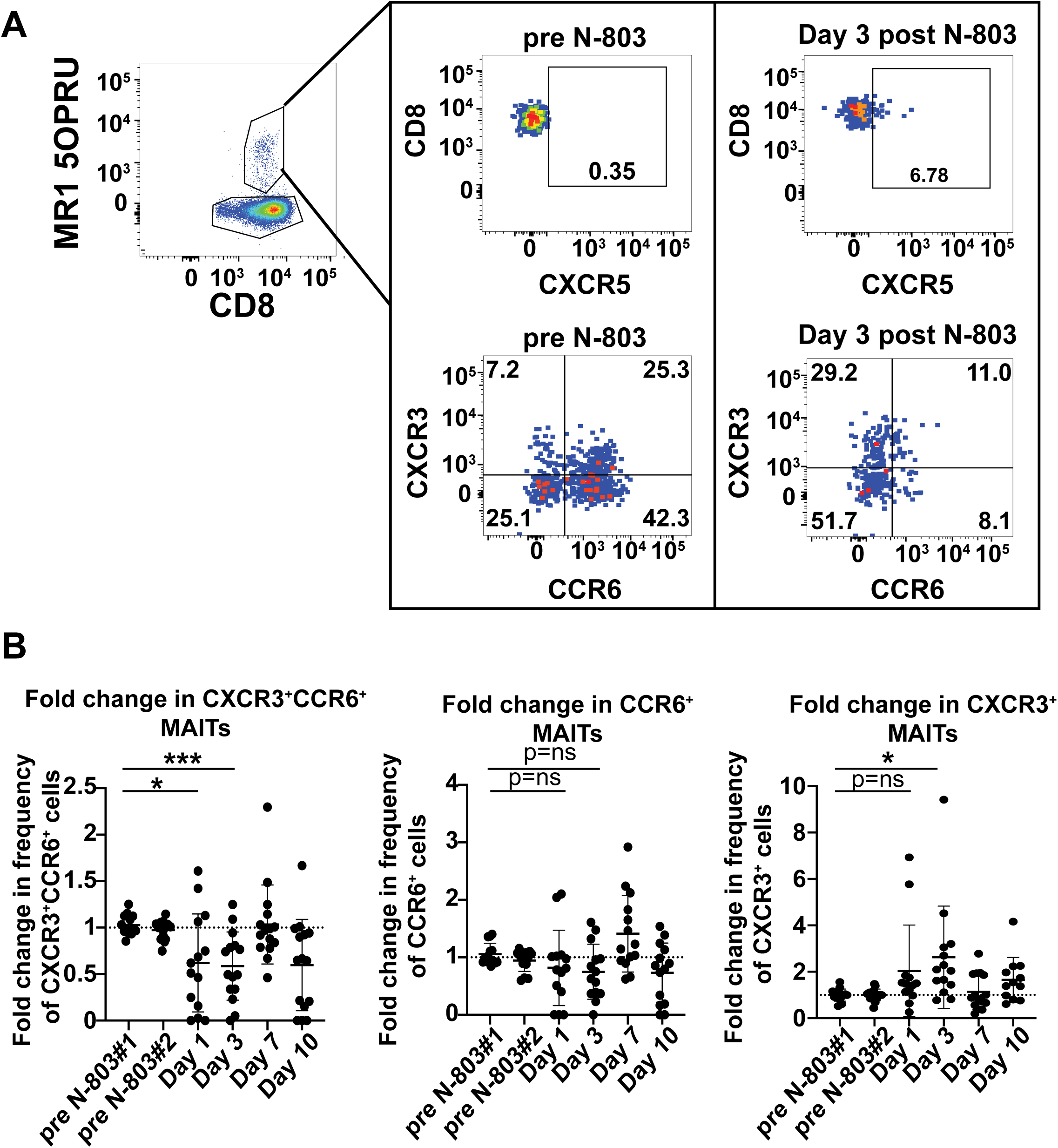
*In vivo* treatment of SIV+ macaques with N-803 decreases the frequency of CXCR3+CCR6+ MAIT cells in the peripheral blood. A, Frozen PBMC collected from the timepoints pre and post N-803 indicated on the axes in (B) were thawed and stained with the panel described in Table 3. Flow cytometry was performed to determine the frequencies of CD8+ MR1 tetramer+ (MAIT cells) expressing chemokine markers CXCR5, CXCR3, and CCR6. Shown is a representative gating schematic for CXCR5, CXCR3, and CCR6 expression on MAIT cells from a pre N-803 treatment timepoint, and 3 days post N-803. B, Cells from the indicated timepoints were stained as described in (A), and the frequencies of MAIT cells that were CXCR3+CCR6+, CCR6+, and CXCR3+ were determined for each timepoint. Then, the frequencies of each subpopulation were normalized relative to the average of the pre N-803 timepoints and graphed as the fold-change in the frequencies of CXCR3+CCR6+ (left), CCR6+ (middle) and CXCR3+ (right) MAIT cells. Repeated measures ANOVA non-parametric tests were performed, with Dunnett’s multiple comparisons for individuals across multiple timepoints. For individuals for which samples from timepoints were missing, mixed-effects ANOVA tests were performed using Geisser-Greenhouse correction. p=ns; not significant; *, p≤0.05; ***, p≤0.0005.

We did not observe any significant differences in the frequencies of CXCR5+ MAIT cells after N-803 treatment compared to pretreatment timepoints (data not shown). We examined the frequencies of CXCR3+, CXCR3+CCR6+, and CCR6+ MAIT cells during N-803 treatment. We normalized the data for each animal relative to the pre N-803 average because there was wide inter-animal variability (Fig. 7B). On days 1 and 3 post N-803 treatment, there was a statistically significant decline in the fold change of CXCR3+CCR6+ MAITs (Fig. 7B, left). The decline in CCR6+ MAIT cells on the same days was not statistically significant (Fig. 7B, middle). There was a corresponding increase in the fold change in CXCR3+ cells on day 3 post N-803 (Fig. 7B, right). Overall, we concluded that N-803 transiently disturbed the population of MAIT cells expressing CXCR3 and CCR6 in the peripheral blood immediately after N-803 treatment.

### N-803 does not alter MAIT cell frequencies in the lymph nodes but increases their ki-67 and Granzyme B expression

N-803 treatment caused a decline in MAIT cells in the peripheral blood (Fig. 5), as well as changes in the frequency of MAIT cells expressing chemokine receptors associated with trafficking to sites of immune activation (Fig. 6). Therefore, we wanted to determine if MAIT cells were trafficking from the peripheral blood to the lymph nodes or tissue sites after N-803 treatment. Lymph node samples were collected on days 1 and 7 post N-803 treatment (Fig 4). MAIT cells were present at lower frequencies in the lymph nodes compared to MAIT cells in the blood or BAL (Supplementary Fig. 2), which was similar to what has been observed in previous studies in SIV-naïve macaques (36). We found that N-803 treatment did not affect MAIT frequencies (Fig. 8A, left). MAIT cells in the lymph nodes had a predominant CM phenotype, and N-803 treatment did not alter the frequencies of CM (Fig. 8A, middle) or EM MAIT cells (Fig. 8A, right).

**Fig. 8.**
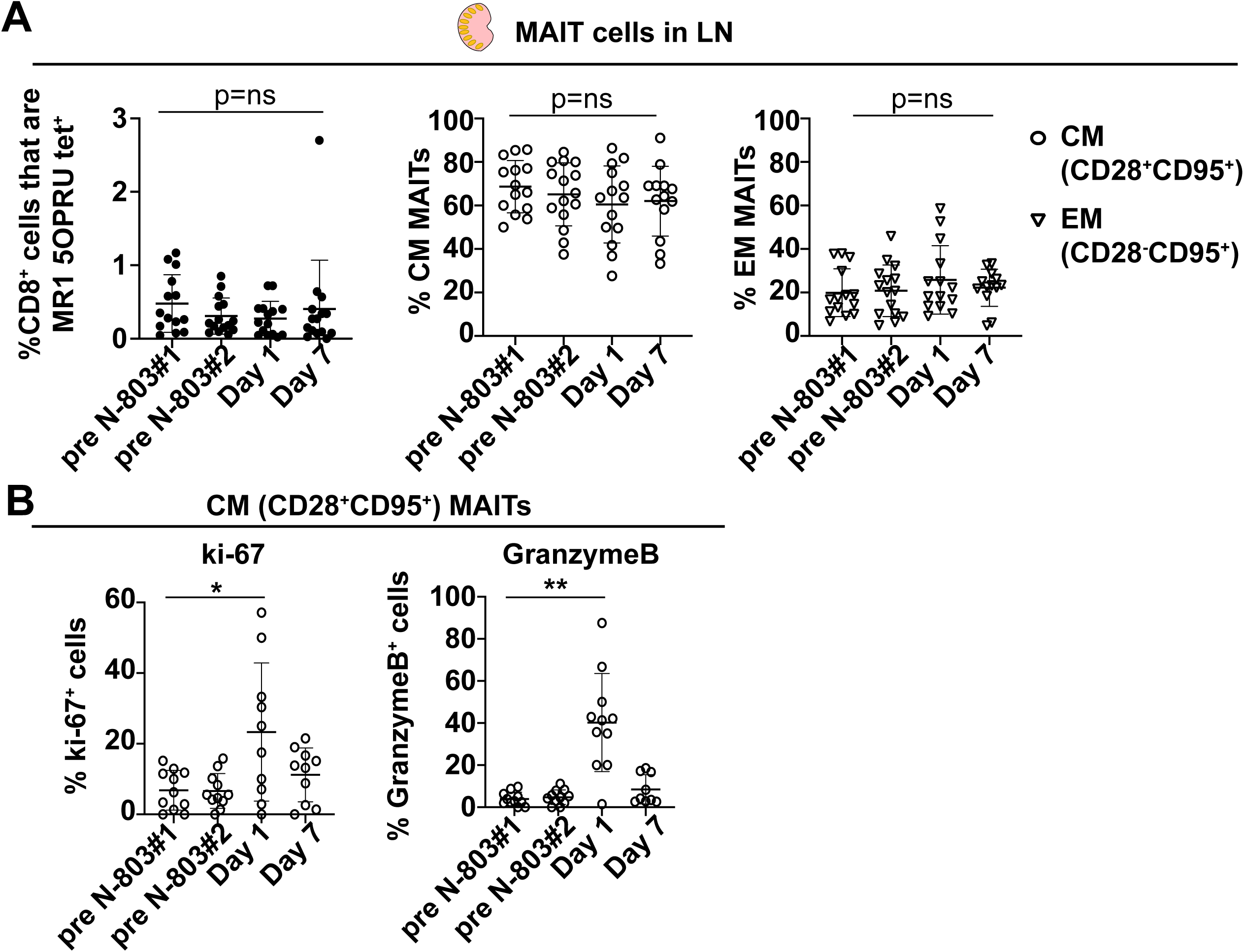
N-803 treatment *in vivo* increases the frequency of ki-67 and Granzyme B+ MAIT cells present in the lymph nodes one day after treatment. A, Frozen cells isolated from lymph node samples that were collected at the indicated timepoints pre and post N-803 treatment were thawed, and flow cytometry was performed with the panel described in Table 2. The frequency of MAIT cells (CD8+ MR1 tet+ cells, left panel), as well as the frequencies of MAIT cells that were central memory (CM; CD28+CD95+, open circles, middle panel) or effector memory (EM; CD28-CD95+, open triangles, right panel) were determined for each timepoint. B, Central memory (CM) MAIT cells in the lymph nodes from the indicated timepoints were stained as in (A), and the frequencies of ki-67+ (left panel) and Granzyme B+ (right panel) cells were determined. Repeated measures ANOVA non-parametric tests were performed, with Dunnett’s multiple comparisons for individuals across multiple timepoints. For individuals for which samples from timepoints were missing, mixed-effects ANOVA tests were performed using Geisser-Greenhouse correction. p=ns, not significant; *, p≤0.05.

We also examined the frequency of CM MAIT cells in the lymph node expressing the proliferation marker ki-67 and the degranulation marker granzyme B on days 1 and 7 after N-803 treatment. There were too few EM MAIT cells for accurate characterization (data not shown). We observed significant increases in the frequencies of ki-67+ and Granzyme B+ CM MAIT cells in the lymph nodes one day after N-803 treatment (Fig. 8B). Similar to the peripheral blood, this effect was transient and returned to pre-treatment levels by 7 days post treatment.

### N-803 affects the phenotype of MAIT cells in the airways

We also assessed the frequency and phenotypes of MAIT cells in the airways during N-803 treatment. We collected BAL fluid prior to, and then 3 days after N-803 treatment (Fig. 4). We found that N-803 treatment did not affect the frequency of MAIT cells (Fig. 9B). N-803 also did not alter the frequencies of total CD3+ cells or conventional CD3+CD8+ T cells (CD3+CD8+MR1 tetramer-cells) in the BAL for the timepoints we examined (Supplementary Fig. 3).

**Fig. 9.**
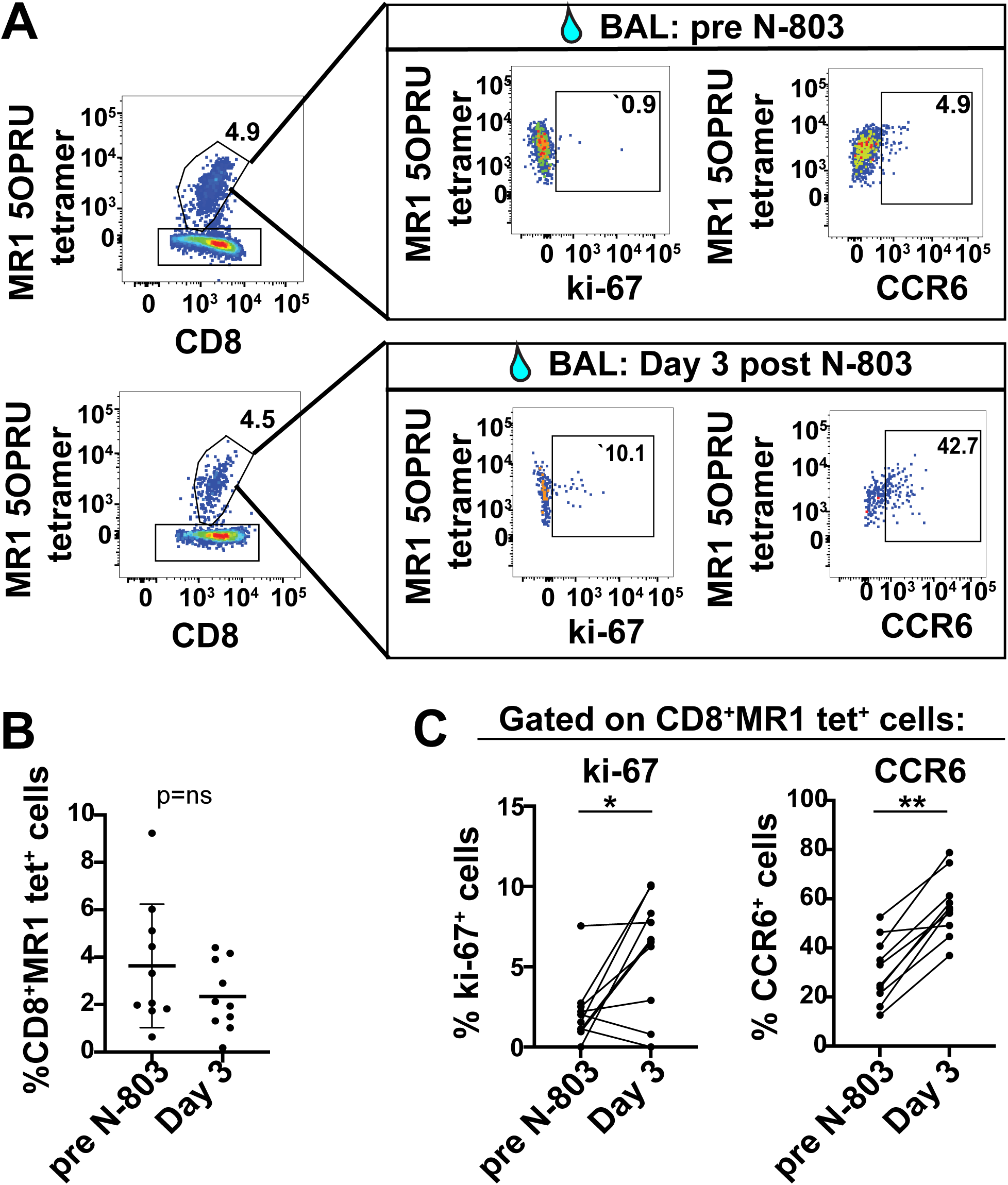
N-803 treatment *in vivo* increases the frequency of ki-67+ and CCR6+ MAIT cells present in the airways. A, Cells isolated from freshly-obtained bronchoalveolar lavage (BAL) from the indicated timepoints pre and post N-803 treatment were stained with the antibodies indicated in Table 4 for flow cytometric analysis. Shown is a representative flow plot of MAIT cells from one individual, as well as the ki-67 and CCR6 expression on those cells, from pre and day 3 post N-803 timepoints. B, The frequencies of MAIT cells (CD8+ MR1 tet+ cells) was determined for the indicated timepoints pre and post N-803 as described in (A). C, The frequency of MAITs expressing ki-67 (left panel) and CCR6 (right panel) were determined as described in (A). Wilcoxon matched pairs rank-signed tests were performed. p=ns, not significant; *, p≤0.05.

We characterized the frequency of MAIT cells expressing activation markers CD69, HLA-DR, and CD154, the proliferation marker ki-67, and the trafficking marker CCR6 in the BAL fluid. An example of typical staining for MAIT cells, as well as ki-67 and CCR6, is shown for BAL cells collected both before and after N-803 treatment in Fig. 9A. There were no changes in the frequencies of CD69, HLA-DR, or CD154+ cells pre or post N-803 treatment (data not shown). There were significant increases in the frequencies of ki-67+ and CCR6+ MAIT cells on day 3 post N-803, relative to the average of the pre N-803 time points (Fig. 9C). The increased frequency of CCR6+ MAIT cells in the BAL was coincident with the decrease in CXCR3+CCR6+ and CCR6+ cells in the peripheral blood on day 3 post N-803 treatment (Fig. 6B).

### MAIT cells from the blood and airways of N-803 treated macaques have increased IFNγ production when stimulated with *E.coli* and *M.smegmatis*

N-803 has been shown in previous studies to increase the ability of NK and CD8 T cells to produce cytokines and cytolytic enzymes, both *in vitro* and *ex vivo* (32, 56). Given that we observed an increase in IFNγ production from MAIT cells pre-incubated with N-803 during antigen stimulation in our i*n vitro* functional assays (Figs. 2-3), we hypothesized that MAIT cells from N-803 treated macaques could also have improved function *ex vivo*. We performed functional assays with total cells collected from PBMC and BAL both before and after N-803 treatment. Too few MAIT cells were present in the LN to perform similar assays (data not shown). Cells were incubated with 10 CFU of either *E.coli* or *M.smegmatis*. The frequencies of IFNγ, TNFα, and CD107a+ MAITs were then determined (Fig. 10).

**Fig. 10.**
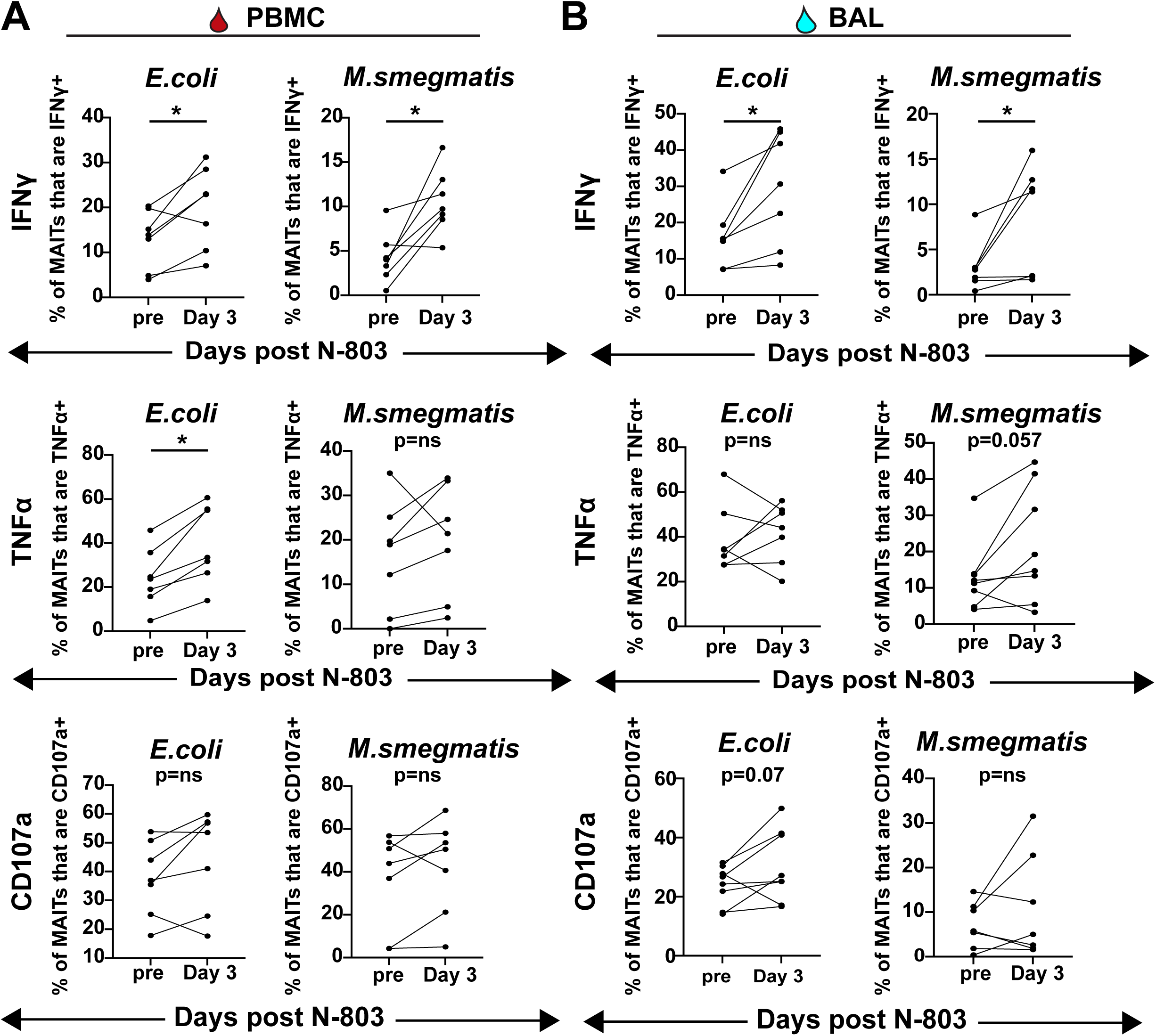
N-803 treatment *in vivo* increases the frequency of IFNγ production from MAIT cells stimulated *ex vivo* with *E.coli* or *M.smegmatis*. A and B, Cells from either PBMC (A) or bronchoalveolar lavage fluid (BAL, B) from pre and day 3 post N-803 timepoints were stimulated either overnight (PBMC, A) or for 6 hours (BAL, B) with 10 colony forming units (CFU)/cell of either *E.coli* or *M.smegmatis*. The frequencies of MAIT cells expressing IFNγ (top), TNFα (middle), or CD107a (bottom) were determined for each timepoint. The data for each stimulus (*E.coli*, left graphs, *M.smegmatis*, right graphs) are shown as the frequency of IFNγ+, TNFα+, or CD107a+ after background subtraction. Wilcoxon matched-pairs Rank signed tests were performed. p=ns, not significant; *, p≤0.05.

We found that the frequency of MAIT cells from the PBMC and BAL producing IFNγ after stimulation with *E.coli* or *M.smegmatis* was higher when using cells collected from animals three days after receiving N-803, when compared to the pre N-803 timepoints (Fig. 10A and B, top panels). While the frequency of MAIT cells producing TNFα and CD107a trended higher on day 3 post N-803 relative to pre N-803 timepoints, results were inconsistent across tissue type and bacterial stimulus (Fig. 10A and B, middle and lower panels). Overall, we concluded that *in vivo* N-803 treatment could improve MAIT cell *ex vivo* function in assays with bacteria, including mycobacteria.

## Discussion

Here, we show for the first time that an IL-15 superagonist, N-803, can affect MAIT cell trafficking, activation, and function *in vivo* in SIV-infected macaques. We found that N-803 could increase the frequencies of ki-67 and granzyme B+ MAIT cells in several compartments, including the lymph nodes and airways (Figs. 6, 8-9). Finally, MAIT cells present in the PBMC and airways from N-803 treated macaques had improved function *ex vivo* against both *E.coli* and *M.smegmatis* stimuli (Fig. 10). Overall, our results establish that N-803 can improve MAIT cell activity and function *in vivo*.

Like recombinant IL-15, N-803 was able to increase the ability of MAIT cells to produce IFNγ in response to bacterial stimulus *in vitro* (Fig. 2) and *ex vivo* (Fig. 10). This data strengthens the findings from other *in vitro* studies suggesting that MAIT cells can become activated and exhibit improved anti-microbial function both directly and indirectly by IL-15 (19, 57). Importantly, we found this to be true for both *E.coli* and *M.smegmatis* (Figs. 2-3, 10), suggesting that N-803 could improve MAIT cell function both *in vitro* and *in vivo* across a broad spectrum of bacteria. MAIT cells have been implicated in the antimicrobial response to *F. tularensis* (58), *S. typhimurium* (59), *K. pneumoniae* (60), or *C.albicans* (61), for example. Therefore, it is possible that N-803 could be utilized in the future to expand and activate MAIT cells *in vivo* to improve their function against these bacteria.

More specifically, our findings could have implications for the use of N-803 in mycobacterial-directed immunotherapies. Increased numbers of CXCR3+CCR6+ cells trafficking to the airways have been shown to correlated with control of *Mycobacterium tuberculosis* (Mtb) infection in animal models (62). CXCR3+CCR6+ cells are associated with a greater degree of Mtb-specific cytokine production (62). We observed that there was a rapid decline in the frequency of MAIT cells in the peripheral blood expressing CXCR3 and/or CCR6 3 days after N-803 treatment (Fig. 7). This was coincident with an increase in the frequency of CCR6+ cells in the airways (BAL; Fig. 9). In addition to this, N-803 improved both the *in vitro* (Figs. 2-3) and *ex vivo* (Fig. 10) ability of MAIT cells to produce IFNγ in response to mycobacterial stimuli. These findings, while very preliminary, are supported by the findings of others that mice treated with a heterodimeric IL-15 superagonist exhibited decreased tumor growth and increased trafficking of CD8 T cells and NK cells to tissue sites in a manner mediated by CXCR3/CXCL9/10 in a colon carcinoma model (63). Overall, the potential for N-803 to drive CXCR3+CCR6+ MAIT cells and other immune cells to the airways and tissue sites of immune activation is intriguing, and future studies could explore this further.

Of great importance for future studies will be to test the effects of N-803 treatment on MAIT cells in Mtb-infected macaques. The very transient nature of the *in vivo* effects of N-803 on MAIT cells (Figs. 5-10) likely rules out the possibility that it could be used by itself in Mtb-directed immunotherapy. However, future studies could investigate whether N-803 boosted MAIT cells could have a positive impact on the outcome of Mtb challenge, most likely as an adjuvant in combination with other MAIT-directed vaccines. For example, Sakai and colleagues have shown that 5-OPRU expanded MAIT cells were able to reduce bacterial burden if expanded during chronic Mtb infection (16). Therefore, one possibility would be to use N-803 as an adjuvant during 5-OPRU mediated MAIT cell expansion to see if this could further reduce bacterial burden. Another possibility would be to use N-803 in combination with BCG vaccination, which is already used to prevent TB disease globally. BCG vaccination can activate MAIT cells *in vivo* (36); therefore, BCG+N-803 treatment could further improve MAIT function against mycobacteria.

The animals utilized for the *in vivo* portions of this study were SIV+ and ART-naïve at the time of N-803 treatment, and most animals had very high viral loads (>10^4^ gag copies/mL plasma viral loads in 12/15 animals; Ellis-Connell et al., J.Immunol, under review). While we did observe improved MAIT cell function during N-803 treatment in these SIV+ macaques (Figs. 6, 8-10), it remains a possibility that SIV infection could have adversely impaired MAIT cell trafficking or function during N-803 treatment relative to what might have been observed in SIV-naïve macaques. We found that in our *in vitro* studies, MAIT cells from both SIV-naïve and SIV+ animals exhibited an N-803 mediated enhancement of IFNγ production. However, only the MAIT cells from healthy animals exhibited improved TNFα production when treated with N-803 (Fig. 2). Whether this reflects differences between healthy and SIV+ animals, or if it is a consequence of small group sizes, is unknown. It would not be surprising if MAIT cells present in healthy versus SIV+ macaques respond differently to N-803. We previously observed a functional impairment of MAIT cells during Mtb and SIV co-infection (14). In a recent study of longitudinal HIV+ samples, MAIT cells exhibited a more innate-like phenotype over time (64). Similar longitudinal SIV studies in pigtailed macaques found that MAIT cells lost expression of T-bet, which can affect IFNγ production (37, 65). SIV+ macaques represent an important cohort of individuals with regards to exploring the role of immunotherapeutic agents to combat Mtb infection, as the immune cells of HIV/SIV+ individuals often have impaired function, resulting in worse TB disease outcomes than healthy individuals (66, 67). Therefore, future studies could focus on treating SIV-naïve animals with N-803 to determine if MAIT cells behave similarly with regards to trafficking and function.

Our study here focused on the role of N-803 treatment on MAIT cell function, but IL-15 agonists have effects on many other immune cell types *in vivo*. Recombinant IL-15 has been used *in vivo* previously in mouse models of Mtb infection, where it was shown that IL-15 treatment protected against Mtb challenge by expanding Mtb-specific T cells (68). N-803 has already been shown to expand virus-specific cells *in vitro* and *in vivo* models of HIV/SIV infection (23); therefore, it is probable that N-803 could improve the conventional T cell response to Mtb infection as well. IL-15 has also been shown *in vitro* to expand dendritic cells and improve their ability to suppress the growth of Mtb in infected macrophages (69). Overall, the use of N-803 as an adjuvant to Mtb-directed immunotherapies could improve the function of several cell types and lead to improved TB disease outcomes.

Overall, our findings here advance the knowledge of how *in vivo* treatment of macaques with N-803 affects MAIT cell function. N-803 improved MAIT cell function against mycobacterial stimuli, and also trafficked cells away from the peripheral blood during *in vivo* treatment. We also present preliminary data that MAIT cells have altered phenotypes and improved function the airways. There is a growing body of evidence that IL-15 agonists could improve the outcome to Mtb infection. Our findings could have important implications for the use of N-803 in anti-Mtb immunotherapy.

## Acknowledgements

We thank staff at the Wisconsin National Primate Resource Center (WNPRC) for excellent veterinary care of the animals involved in this study.

This study was supported by funding supplied through the National Institute of Health (NIH R01 grant number AI108415).

This research was conducted at a facility constructed with support from Research Facilities Improvement Program grant numbers RR15459-01 and RR020141-01. The Wisconsin National Primate Research Center is also supported by grants P51RR000167 and P51OD011106.

## SUPPLEMENTARY FIGURE LEGENDS

**Supplementary Fig. 1**. Functional assays were performed as indicated in Figs. 2-3, 10 and the methods using PBMC and BAL samples collected prior to and after N-803 treatment. Flow cytometry was performed using antibodies in Tables 6 (PBMC) and 7 (BAL) to measure the frequency of MAIT cells expressing IFNγ, TNFα, or CD107a after stimulation. Shown is a representative flow plot for the frequencies IFNγ, TNFα, or CD107a+ MAIT cells in PBMC stimulated with media control (no stim) or 10 CFU of fixed *E.coli*.

**Supplementary Fig. 2**. A, Cells from BAL (left), PBMC (middle), and lymph node (LN; right) were stained for flow cytometric analysis using panels described in Table 4 (BAL) and Table 2 (PBMC, LN). Shown are representative images of the frequency of CD8+MR1 tetramer+ (MAIT) cells for each tissue type. B, Cells from BAL (blue), PBMC (red), and LN (black) were stained as described in (A). Shown are the average frequencies of CD8+MR1 tetramer+ cells prior to N-803 treatment for all animals described in Fig. 4.

**Supplementary Fig. 3**. A, BAL collected from timepoints indicated in Fig. 4 were stained for flow cytometric analysis using the panel described in Table 4. Shown is a representative gating schematic. B, BAL samples collected from macaques prior to and 3 days after N-803 treatment were stained for flow cytometric analysis as described in (A). The frequencies of CD3+ cells (left) and CD3+CD8+ cells (right) are shown pre and post N-803. Wilcoxon matched-pairs Rank signed tests were performed; p=ns, not significant.

